# Characterization of human iPSC-derived astrocytes with potential for disease modeling and drug discovery

**DOI:** 10.1101/2019.12.28.890012

**Authors:** Vincent Soubannier, Gilles Maussion, Mathilde Chaineau, Veronika Sigutova, Guy Rouleau, Thomas Durcan, Stefano Stifani

## Abstract

Astrocytes play a number of key functions in health and disease. Reactivation of astrocytes occurs in most, if not all, neurological diseases. Most current information on the mechanisms underlying astrogliosis derives from studies using rodent experimental systems, mainly because the ability to study human astrocytes under healthy and pathological conditions has been hampered by the difficulty in obtaining primary human astrocytes. Here we describe robust and reliable derivation of astrocytes from human induced pluripotent stem cells (iPSCs). Phenotypically characterized human iPSC-derived astrocytes exhibit typical traits of physiological astrocytes, including spontaneous and induced calcium transients. Moreover, human iPSC-derived astrocytes respond to stimulation with a pro-inflammatory combination of tumor necrosis factor alpha, interleukin 1-alpha, and complement component C1q by undergoing changes in gene expression patterns suggesting acquisition of a reactive astrocyte phenotype. Together, these findings provide evidence suggesting that human iPSC-derived astrocytes are a suitable experimental model system to study astrocyte function and reactivation in healthy and pathological conditions of the human nervous system.

## 1. INTRODUCTION

Astrocytes play a number of important roles in the healthy nervous system by supporting neuronal activity in several ways, including regulation of synaptic transmission, release of neurotrophic factors, promotion of neuronal metabolic functions, and cross-talk with other glial cells, to name only a few [reviewed by Zhou *et al.*, 2019; Zuchero and Barres, 2015]. Astrocytes are also important players in the injured or diseased nervous system, when they can acquire ‘reactive’ phenotypes, characterized by morphological and gene expression changes during a process often termed astrogliosis [reviewed by Liddelow and Barres, 2017; Siracusa *et al.*, 2019; Valori *et al.*, 2019]. Reactive astrocytes mediate a variety of mechanisms, including release of cytokines and chemokines that can participate in complex neuroinflammatory processes, intercellular communication with microglia involved in immune responses, and modulation of immune cell activation at damaged sites. Reactive astrocytes are thought to contribute to either neuroprotective or neuroinflammatory responses depending on the local microenvironment, although evidence exists that astrogliosis is often a complex process characterized by a combination of both inflammatory and protective effects depending on the stage of the damage or injury process [Li *et al.*, 2019; Liddelow and Barres, 2017; Yamanaka and Komine, 2018].

Astrocyte reactivation is a common event in virtually all neurological diseases [Li *et al.*, 2019; Liddelow and Barres, 2017; Siracusa *et al.*, 2019; Yamanaka and Komine, 2018]. For instance, hallmarks of astrocyte activation feature prominently in *post-mortem* tissues from amyotrophic lateral sclerosis (ALS) patients, as well as in the spinal cord and cranial motor nuclei in ALS transgenic mouse models [Anneser *et al.*, 2004; Baron *et al.*, 2005; Evans *et al.*, 2014; Ferraiuolo *et al.*, 2007; Poloni *et al.*, 2000; Schiffer *et al.*, 1996]. More importantly, increasing evidence suggests that reactive astrocytes contribute directly to the degeneration of motor circuits in ALS animal models by promoting non-cell autonomous mechanisms of motor neuron degeneration [Diaz-Amarilla *et al.*, 2011; Ferraiuolo, 2014; Haidet-Phillips *et al.*, 2011; Ouali Alami *et al.*, 2018; Radford *et al.*, 2015; Yamanaka and Komine, 2018].

Progress in understanding how astrocytes contribute to human neurological diseases has been slow, at least in part due to the fact that most information on the mechanisms underlying astrogliosis has emerged from studies with rodent experimental model systems. There is evidence that rodent and human astrocytes exhibit a number of gene expression and functional differences suggesting that rodent astrocytes may not always represent informative model systems for human astrocytes, especially in neurodegenerative conditions [Oberheim *et al.*, 2009; Tarassishin *et al.*, 2014; Zhang *et al.*, 2016]. Moreover, the paucity of studies based on human primary astrocytes has limited the ability to study human astrocytes carrying the genetic alterations found in familial form of neurodegenerative diseases, thus affecting the goal of complementing studies using experimental animal models with human cell-based models. This situation has recently started to change due to progress in the application of induced pluripotent stem cell (iPSC) technologies in the generation of human iPSC-derived astrocytes [Serio *et al.*, 2013; Shaltouki *et al.*, 2013; Tcw *et al.*, 2017]. This remarkable progress offers previously unavailable experimental opportunities to study human astrocyte biology, model astrocyte pathophysiological mechanisms *in vitro,* and design large-scale screens of new potential therapeutic compounds targeting astrocytes.

Here we describe the generation and phenotypic characterization of astrocytes derived from human iPSCs, and show that these human iPSC-derived astrocytes display molecular and biological features typical of physiological astrocytes. In particular, human iPSC-derived astrocytes respond to astrogliosis-inducing conditions by undergoing changes in gene expression patterns suggestive of a transition to a reactive phenotype. These findings suggest that human iPSC-derived astrocytes represent physiologically-informative experimental model systems to study astrocyte function and reactivation in healthy and diseased conditions of the human nervous system.

## 2. METHODS

### 2.1. Human iPSC lines

Human iPSC line NCRM-1 (male) was obtained from the National Institutes of Health Stem Cell Resource (Bethesda, MD, USA). AJG-001-C4 control iPSC line was established from peripheral blood mononuclear cells (PBMC) obtained from a 37-year-old male through episomal reprogramming. AIW-002-02 iPSC line was established from PBMC obtained from a 37-year-old male through retrovirus reprogramming. These procedures were conducted under Ethical Review Board approval by McGill University. Both AJG 001-C4 and AIW-002-2 iPSC lines were generated at the Montreal Neurological Institute through the Open Biorepository, C-BIGR, under an Open Science framework. Pluripotent state of human iPSC lines was routinely assessed by testing for expression of the stem cell markers NANOG and OCT4.

### 2.2. Derivation of neural progenitor cells from human iPSCs

Human iPSCs at low passage number were cultured in mTeSR medium (STEMCELL Technologies; Vancouver, BC, Canada; Cat. No. 85850) in 10-cm culture dishes (Thermo-Fisher Scientific; Waltham, MA, USA; Cat. No. 353003) coated with Matrigel (Thermo-Fisher Scientific; Waltham, MA, USA; Cat. No. 08-774-552) until they reached 70%-80% confluence. At this time point (designated as day *in vitro* 0 - DIV0), the culture medium was removed, cells were rinsed with phosphate-buffered saline (PBS) (5 ml) and detached by addition of ‘Gentle Dissociation Reagent’ (3 ml) (STEMCELL Technologies; Cat. No. 07174) stored at room temperature. After 3-5 minutes in Gentle Dissociation Reagent, cells were recovered and transferred to a 15-ml Falcon tube (Corning; Corning, NY, USA; Cat. No. 352070). Dulbecco’s minimum essential medium (DMEM)/F12 medium (Life Technologies; 10565-018) was added (7 ml), followed by centrifugation at 1,300 rpm for 3 min. Cell pellets were resuspended in mTeSR medium (1 ml) and transferred into 6-well plates coated with Matrigel. Cells were seeded at 15%-25% confluence and cultured in mTeSR medium (2 ml) in the presence of the Rock inhibitor Y-27632 (Stemgent; Cambridge, MA, USA; Cat. No. 04-0012) at 37°C. The next day (designated DIV1), culture medium was replaced with 2.5 ml of ‘Neural Induction Medium 1’ [DMEM/F12 medium supplemented with 1% N2 (Life Technologies; Cat. No. 17504-044), 2% B27 (Life Technologies; Cat. No. 17502-048), 0.1% bovine serum albumin (BSA) (Life Technologies; Cat. No. 15260-037), 2 μM SB431542 reagent (Tocris Bioscience; Bristol, UK; Cat. No. 1614), 200 ng/ml Noggin (PeproTech; Rocky Hill, NJ, USA; Cat. No. 120-10c), 1 mM GlutaMax (Life Technologies; Cat. No. 35050-061), 1% non-essential amino acids (Life Technologies; Cat. No. 11140-050), 1 μg/ml laminin (Sigma Aldrich; St. Louis, MO, USA; Cat. No. L2020), 50 U/ml penicillin G, 50 mg/ml streptomycin (Life Technologies; Cat. No. 15140-122)]. This culture medium was changed (2.5 ml) every two days and cells were cultured until DIV4, when cells received fresh medium (5 ml) and were cultured until they reached confluence (usually by DIV7).

At DIV7, Neural Induction Medium 1 was replaced with 5 ml of ‘Neural Induction Medium 2’ (DMEM/F12 medium supplemented with 1% N2, 2% B27, 0.1% BSA, 1 mM GlutaMax, 1% nonessential amino acids, 1 μg/ml laminin, 50 U/ml penicillin G, 50 mg/ml streptomycin). Medium was replaced every day until DIV12, when cells were dissociated by incubation with Accutase (Innovative Cell Technologies; San Diego, CA, USA; Cat. No. AT-104) at 37 ^o^C for 3-5 min and transferred to either Petri dishes or ‘low attachment’ dishes (Fisher Scientific; Cat. No. FB0875712), followed by culture in ‘Neural Progenitor Cell (NPC) Expansion Medium’ [DMEM/F12 medium supplemented with 1% N2, 2% B27, 20 ng/ml EGF (PeproTech; Cat. No. AF-100-15), 20 ng/ml FGF2 (PeproTech; Cat. No. 450-18b), 1 mM GlutaMax, 1% non-essential amino acids, 1 μg/ml laminin, 50 U/ml penicillin G, 50 mg/ml streptomycin] in the presence of Rock inhibitor Y-27632. Cells were then cultured until formation of free-floating cellular spheres (‘neurospheres’), which usually occurred in 2 to 3 days.

In each differentiation procedure, neurospheres were collected using a 40 μm Nylon mesh cell strainer (Corning; Cat. No. 352340) placed over a 50-ml Falcon tube. Cells recovered on cell strainer were rinsed with PBS (5 ml) and then transferred to a 10-cm Matrigel-coated culture dish by flushing with 10 ml NPC Expansion Medium. Another cycle of neurosphere formation and isolation was usually performed by repeating the same steps in order to increase the number of NPCs obtained. On average, ~90% of induced cells were positive for the expression of NPC markers SOX1, SOX2, and PAX6. NPCs obtained following this protocol can be frozen and stored in liquid nitrogen, with usually good cell survival rates upon thawing.

### 2.3. Differentiation of astrocytes from human iPSC-derived neural progenitor cells

Induced human NPCs were differentiated into astrocytes essentially as described previously [Tcw *et al.*, 2017], with minor modifications. In each derivation, NPCs were seeded at low cell density (15,000 cells/cm_2_) in two T25 flasks in the presence of 5 ml of NPC expansion medium containing Rock inhibitor. The following day, medium was replaced with ‘Astrocyte Differentiation Medium 1’ [ScienCell Astrocyte Growth Medium (ScienCell Research Laboratories; Carlsbad, CA, USA; Cat. No. 1801b), astrocyte growth supplement (ScienCell Research Laboratories; Cat. No. 1852), 2% fetal bovine serum (FBS) (ScienCell Research Laboratories; Cat. No. 0010), 50 U/ml penicillin G, 50 mg/ml streptomycin]. Half medium was replaced with fresh medium every 3 to 4 days and cells were maintained under these culture conditions for 30 days without passaging. On average, at least 75% of induced cells were positive for the expression of astrocytes markers, such as S100beta (S100β), connexin 43, and aquaporin 4 (AQ4) after 30 days. At this stage, induced astrocytes were proliferative and were either cultured further or frozen and stored in liquid nitrogen for future use. For the purpose of continued culture, induced astrocytes were switched to ‘Astrocyte Differentiation Medium 2’ (same as Astrocyte Differentiation Medium 1 but lacking FBS) after 30 days and then cultured under these conditions for 2-3 months, depending on the experimental endpoints. Induced astrocytes were routinely examined for the expression of GFAP, S100β, and AQ4.

### 2.4. Characterization of human iPSC-derived cells by immunocytochemistry

Cultured human iPSCs, induced NPCs and astrocytes were analyzed by immunocytochemistry, which was performed as described previously [Methot *et al.*, 2018]. The following primary antibodies were used: rabbit anti-NANOG (1:500; Abcam; Cambridge, UK; Cat. No. Ab21624); goat anti-OCT3/4 (1:500; Santa Cruz Biotechnology; Dallas, TX; Cat. No. Sc-8628); goat anti-SOX1 (1/500; R&D Systems; Minneapolis, MN, USA; Cat. No. AF3369); goat anti-SOX2 (1/500; R&D Systems, Cat. No. AF2018); rabbit anti-PAX6 (1/500; Covance; Emeryville, CA, USA; Cat. No. PRB-278P); mouse anti-NESTIN (1/500; Millipore; Cat. No. MAB5326); mouse anti-glial fibrillary acid protein (GFAP) (1/1,000; Sigma-Aldrich, St. Louis, MO, USA; Cat. No. G3893); mouse anti-S100β (1/1,000; Sigma-Aldrich; Cat. No. S2532); rabbit anti-AQ4 (1:500; Sigma-Aldrich; Cat. No. HPA014784). Secondary antibodies against primary reagents raised in various species were conjugated to Alexa Fluor 555, Alexa Fluor 488, or Alexa Fluor 350 (1/1,000, Invitrogen; Burlington, ON, Canada). Images were acquired with a Zeiss Axio Observer Z1 Inverted Microscope using 20X magnification and a ZEISS Axiocam 506 mono camera.

### 2.5. Detection of calcium transients in human iPSC-derived astrocytes

Sixty days after the start of the differentiation protocol, human iPSC-derived astrocytes were incubated with the calcium indicator Fluo-4 AM (1μM; ThermoFisher Scientific; Carlsbad, CA; Cat. No. F14201) for 30 minutes, followed by two rinsing steps with ScienCell Astrocyte Growth Medium (Cat. No. 1801b), supplemented with antibiotics, for 10 min each. Time-lapse microscopy was then performed at 0.5 Hz with a Zeiss Axio Observer Z1 Inverted Microscope using 20X magnification and a ZEISS Axiocam 506 mono camera. Temperature (37°C), humidity and CO2 (5%) were maintained constant throughout the course of the experiments through TempModule S and CO2 moduleS devices (PeCon GmbH; Erbach, Germany). LED 488 nm was set at 20% and exposure time was 200 ms to avoid phototoxicity. For spontaneous calcium wave acquisition, recording was conducted for 5 min and focus stabilisation was achieved with the use of Zeiss Definite focus every 30 images. For induced calcium wave acquisition, recording was performed for 1 min, before the addition of either 10μM glutamate or 10μM ATP, followed by recording for an additional 4 min.

### 2.6. Treatment of human iPSC-derived astrocytes with TNFα, IL-1α, and C1q

Sixty days after the start of the differentiation protocol, human iPSC-derived astrocytes were incubated with tumour necrosis factor alpha (TNFa) (30 ng/ml; Cell Sciences; Newburyport, MA; Cat. No. CRT100B), interleukin alpha (IL-1a) (3 ng/ml; Sigma-Aldrich; Cat. No. I3901-5UG) and complement component 1, subcomponent q (C1q) (400 ng/ml; MyBioSource; San Diego, CA; Cat. No. MBS143105) for 24 hours, followed by RNA extraction and cDNA synthesis. Each cell pellet was lysed in QIAzol Lysis Reagent (Qiagen; Hilden, Germany; Cat. No. 79306) and total RNA extracted with the miRNEasy Kit (Qiagen; Cat. No. 217004) following the instructions provided by the manufacturer. Reverse transcription was performed on total RNA fractions to obtain cDNA in 40 μl volume containing 400 ng of total RNA, 0.5 μg oligoDT(16) primers, 0.5 mM dNTPs, 0.01 M DTT, and 400 U M-MLV reverse transcriptase (ThermoFisher Scientific; Cat. No. 28025013). Real-time PCR experiments were performed using a QuantStudio 3 Real Time PCR system (ThermoFisher Scientific) in a 10 μl total reaction volume containing 5 μl of fast advanced master mix, 1 μl 20X taqman assay (Thermofisher Scientific), 1 μl of cDNA and 3 μl of RNAse-free water. Analysis of gene expression was conducted using the following oligonucleotide primers: Taqman probes *SERGLYCIN, Hs01004159_m1; IL-1beta, Hs01555410_m1; TNF-alpha, Hs 00174128_m1; SERPING1, Hs00163781_m1, GLUTAMINE SYNTHETASE (GLUL), Hs00365928_g1; SLC1A3/EAAT1/GLAST, Hs00904823_g1; VIMENTIN, Hs00958111_m1; GFAP, Hs00909233_m1; ACTB Hs01060665_g1; GAPDH Hs02786624_g1).* Primers/probesets used were obtained from ThermoFisher Scientific. Data were normalized with *beta-actin* and *GAPDH.* Relative quantification (RQ) was estimated according to the ΔΔCt method [Schmittgen and Livak, 2008].

## 3. RESULTS

### 3.1. *In vitro* generation and molecular characterization of neural progenitor cells from human iPSCs

Neural induction of human iPSCs (three different lines were used) was stimulated by inhibiting TGF-β/BMP-4 signaling using SB431542 compound in combination with Noggin as described in the Methods section. Neuralized cells were tested for the expression of NPC markers, including NESTIN, SOX1, SOX2, and PAX6, 7-14 days after the start of neural induction (DIV 7-14). The majority of neuralized cells expressed these NPC markers at DIV14 (Figure 1). In contrast, little or no expression of the stem cell markers NANOG and OCT-4 was detected at DIV14 (Supplemental Figure S1). These results provide evidence of robust generation of induced cells with features of NPCs from the different human iPSC lines used in these *in vitro* studies.

**Figure 1.**
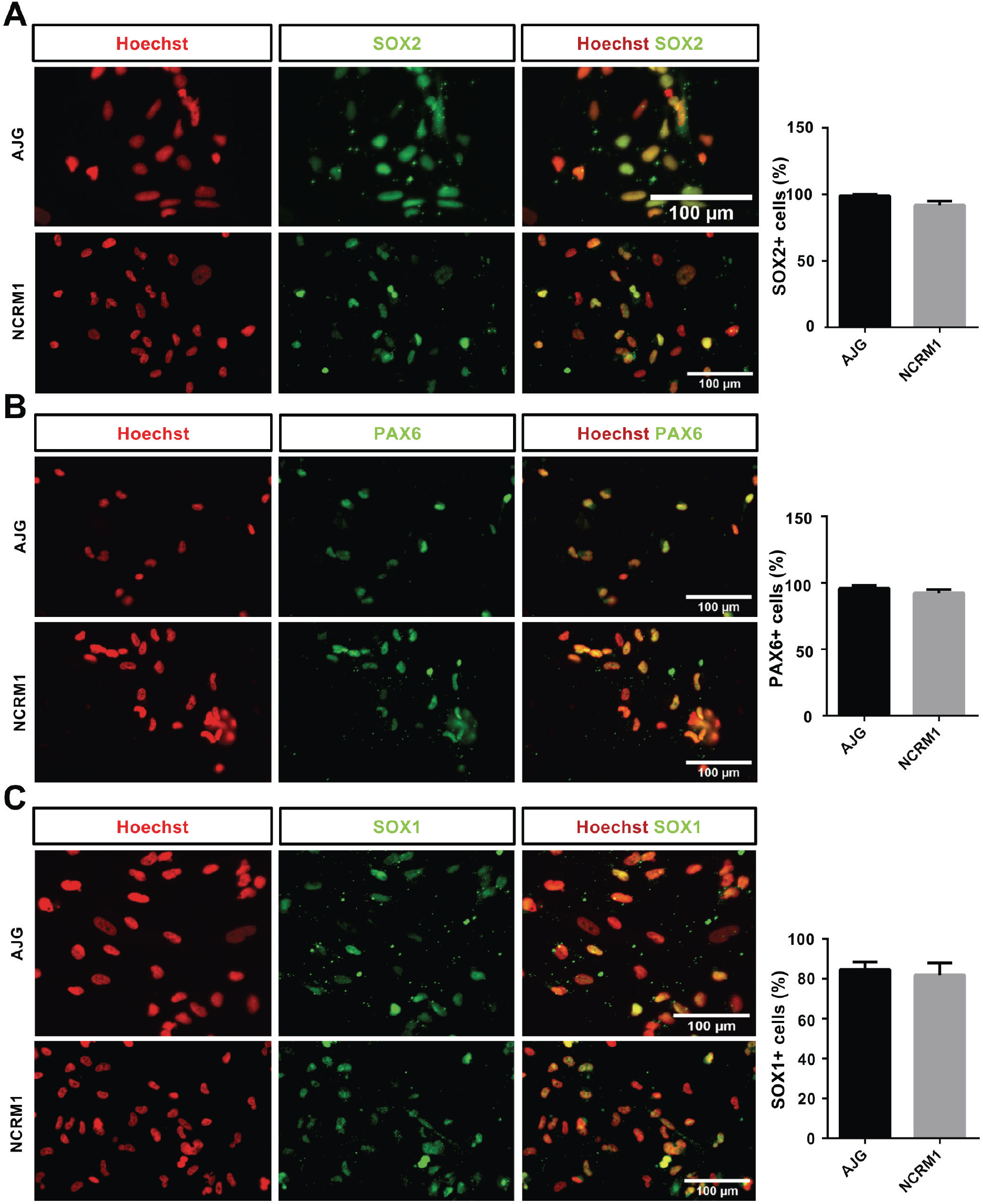
Characterization of human iPSC-derived neural progenitor cells. Representative immunofluorescence analysis of either SOX2, PAX6, or SOX1 expression, as indicated, in NPCs derived from two different iPSC lines (AJG and NCRM1). Cells were counterstained with Hoechst reagent. Quantifications of the fraction of NPCs expressing SOX2, PAX6 or SOX1 are shown in the right-hand column (means ± SEM; n = 3 separate cultures).

### 3.2. *In vitro* differentiation and molecular characterization of astrocytes from human iPSCs

Astroglial differentiation of human iPSC-derived NPCs was promoted by exposure to a defined astrocyte medium from ScienCell, as previously described [Tcw *et al.*, 2017], and as detailed in the Methods section. Under these culture conditions, induced cells exhibiting morphological and immunological features of astrocytes were already detectable at DIV30 (Figure S2). At DIV60, many induced cells (~70%-75% on average) were positive for the expression of both S100β and GFAP, two classical markers of astrocyte identity (Figure 2A). Induced cells also expressed additional astrocyte-enriched proteins/genes, including the gap junction protein connexin 43 (Figure 2B), and the genes *VIMENTIN, GLUTAMINE SYNTHETASE (GLUL),* and *EXCITATORY AMINO ACID TRANSPORTER 1 (EAAT1/ SLC1A3/GLAST)* (Figure 2C). Expression levels of these genes varied between astrocytes derived from different human iPSC lines (cf. Figures 2 and 4 below). At DIV90, human iPSC-derived astroglial cells exhibited mainly flattened morphologies, which included star-shaped cells (Figure 2D). Together, these findings show robust generation of induced cells from human iPSCs with molecular and morphological traits typical of astrocytes *in vitro*.

**Figure 2.**
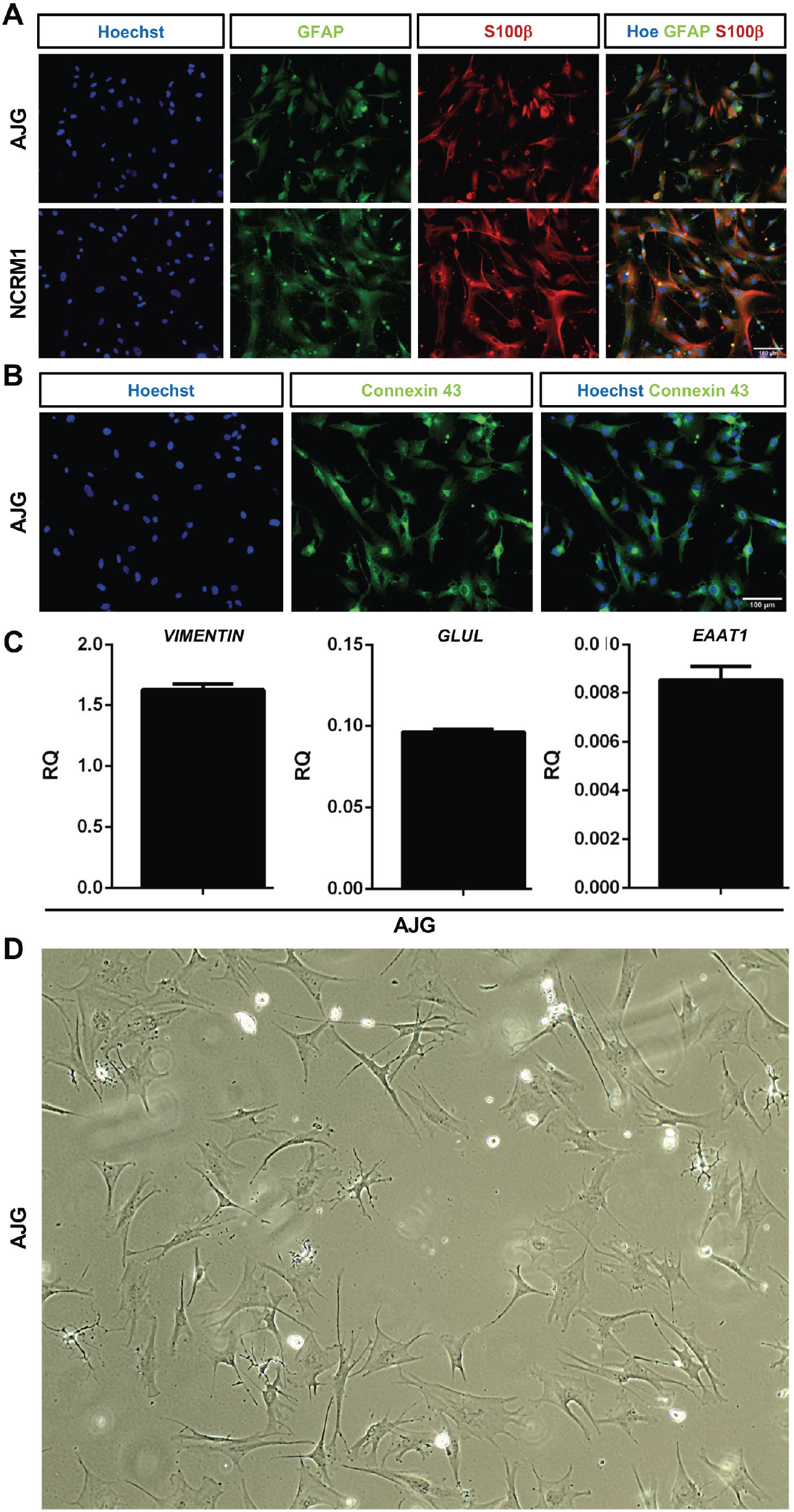
Characterization of human iPSC-derived astrocytes. (A) Representative doublelabeling immunofluorescence analysis of GFAP and S100β expression in DIV60 astrocytes derived from two different iPSC lines (AJG and NCRM1). Cells were counterstained with Hoechst reagent. (B) Immunofluorescence analysis of connexin 43 expression in DIV60 astrocytes derived from iPSC line AJG. Cells were counterstained with Hoechst reagent. (C) Real-time PCR analysis of the expression of *VIMENTIN, GLUL* and *EAAT1* in DIV60 astrocytes derived from human iPSC line AJG. (D) Phase contrast image of DIV90 astrocytes derived from human iPSC line AJG showing presence of cells with flattened morphologies including stellate-like cells.

### 3.3. Human iPSC-derived astrocytes display spontaneous, ATP-induced and glutamate-induced calcium waves

Cytoplasmic Ca^2+^ level fluctuations represent a key component of astrocytic intercellular signaling [Shigetomi *et al.*, 2016]. To determine whether human iPSC-derived astrocytes would exhibit spontaneous and/or induced calcium transients, DIV60 astrocytes were incubated with the Ca^2+^ indicator, Fluo-4 AM, in the absence (spontaneous Ca^2+^ transients) or presence of pulses of extracellular glutamate (10 μM) or ATP (10 μM) (induced Ca^2+^ transients). Human iPSC-derived astrocytes displayed spontaneous calcium spikes, suggestive of intercellular signaling among interconnected cells (Figure 3A). Moreover, a single application of either glutamate or ATP resulted in calcium wave responses (Figures 3B, C). Together, these results provide evidence that human iPSC-derived astrocytes display calcium transients *in vitro* similar to those exhibited by physiological astrocytes, suggesting further that human iPSCs can serve as sources of biologically-relevant human astrocytes.

**Figure 3.**
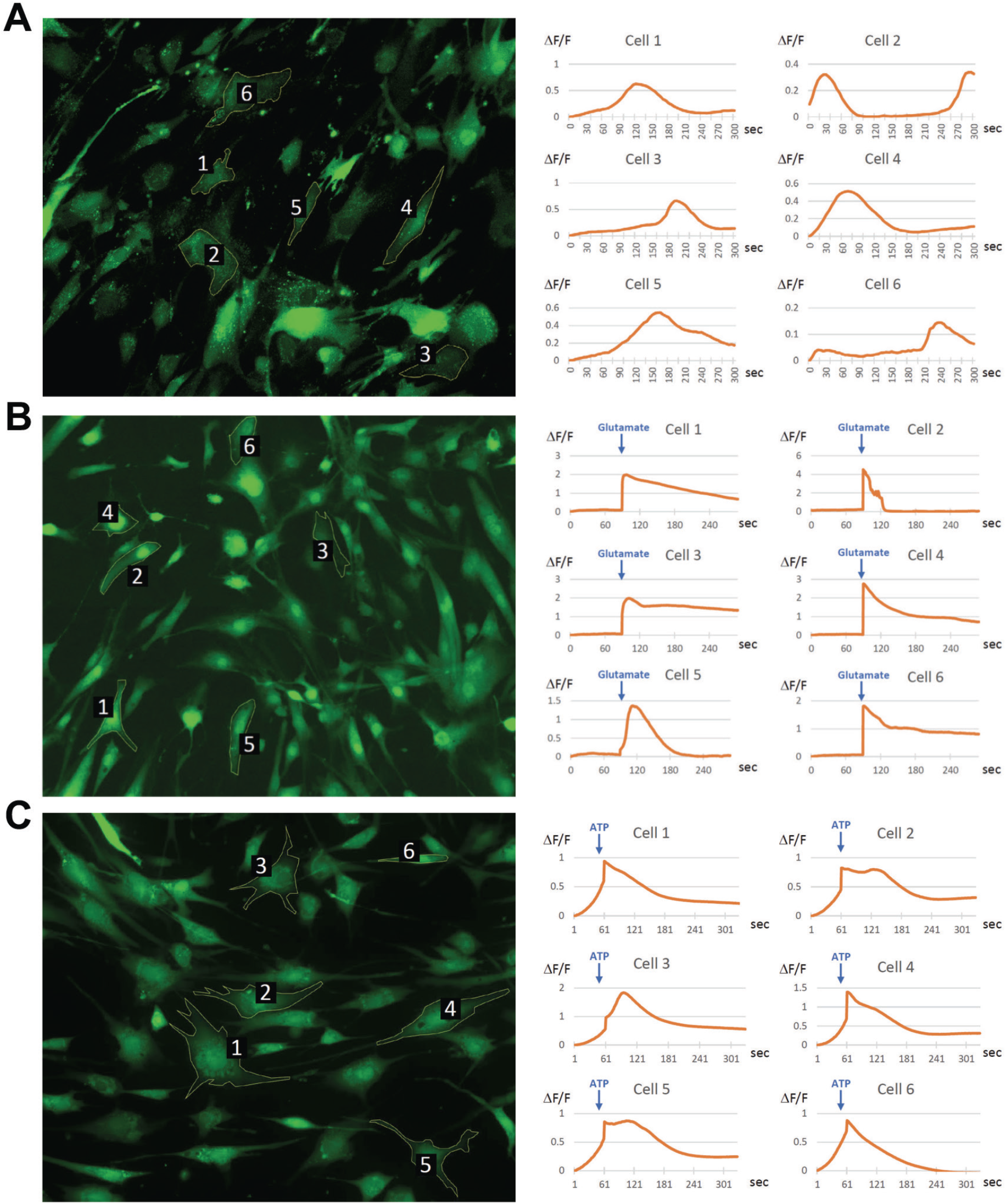
Spontaneous and induced calcium transients in human iPSC-derived astrocytes. (A) Representative Fluo-4 AM visualized DIV60 astrocytes derived from iPSC line NCRM1 exhibiting spontaneous calcium waves. (B, C) Representative glutamate-(B) or ATP-responsive (C) calcium transients in human iPSC-derived astrocytes. Graphs showing immunofluorescence (ΔF/F) over time for 6 randomly selected cells in each case are shown on the right-hand side.

### 3.4. Human iPSC-derived astrocytes acquire a reactive phenotype after exposure to tumor necrosis factor alpha, interleukin 1-alpha and complement C1q

Previous studies have shown that primary rodent brain astrocytes respond to exposure to a combination of TNFa, IL-1a, and C1q by upregulating the expression of several genes associated with an ‘A1-type’ reactive astrocyte phenotype [Clarke *et al.*, 2018; Liddelow *et al.*, 2017]. To determine whether human iPSC-derived astrocytes would respond in a similar manner to treatment with TNFa, IL-1a and C1q, we exposed DIV60 astrocytes to these factors for 24 hours, followed by analysis of gene expression by real time RT-PCR. These studies showed that transcripts previously associated with an A1-type reactive astrocyte phenotype in rodent cells, such as *SERGLYCIN* and *SERPING1*, were upregulated in human iPSC-derived astrocytes (Figure 4A, B). Moreover, treatment with TNFa, IL-1a, and C1q led to increased levels of two genes, *EAAT1* and *GLUL,* previously shown to be induced in reactive astrocytes (Bellaver *et al.*, 2017; Beschorner *et al.*, 2007) (Fig. 4C-D). Consistent with these findings, treated human iPSC-derived astrocytes also displayed increased levels of neuroinflammatory cytokine-encoding transcripts, such as *TNFa* and *IL-1β* (Figure 4E, F). Based on these results, we examined further whether TNFa/IL-1a/C1q-stimulated human iPSC-derived astrocytes would display a pro-inflammatory phenotype by monitoring the levels of transcripts encoding intermediate filament proteins, such as *GFAP* and *VIMENTIN*, which are usually upregulated in pro-inflammatory reactive astrocytes (Clarke *et al.*, 2018; Liddelow *et al.*, 2017; Zamanian *et al.*, 2012). Unexpectedly, neither of these transcripts was increased, rather levels were either decreased or not obviously affected (Figure 4G, H). Similar results were obtained in astrocytes derived from different human iPSC lines (Figure S3). Taken together, these findings provide evidence that human iPSC-induced astrocytes respond to exposure to inflammatory conditions by acquiring a mixed reactive phenotype. These results in turn imply that human iPSC-derived astrocytes may provide an informative experimental model system to study the roles of astrocytes in health and disease.

**Figure 4.**
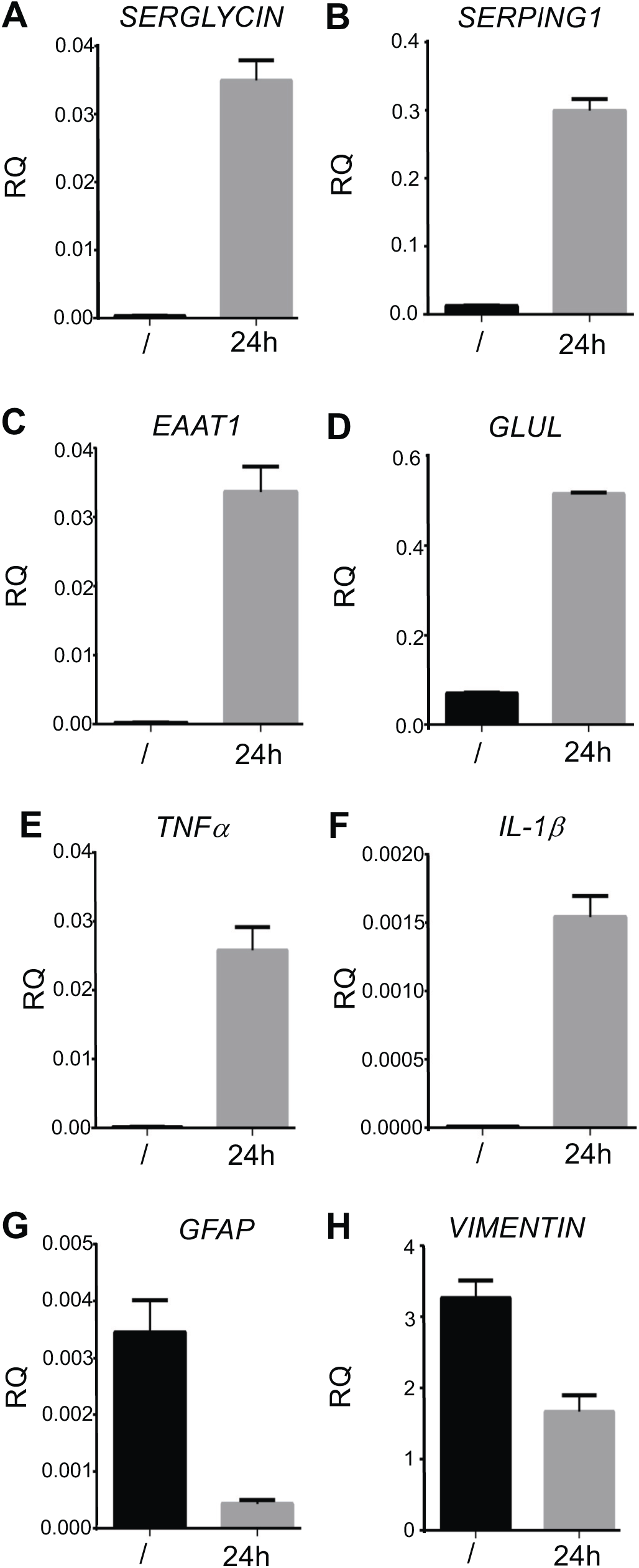
Response of human iPSC-derived astrocytes to TNFα, IL-1α and C1q. Results of real-time PCR experiments depicting mRNA levels of *SERGLYCIN, SERPING1, EAAT1, GLUL, TNFa, IL-1 β, GFAP,* and *VIMENTIN* in DIV60 astrocytes derived from iPSC line AIW not treated (/) or treated for 24h (24h) with a combination of TNFa, IL-1a and C1q. n = 2 experiments performed in triplicates.

## 4. DISCUSSION

Numerous studies are providing increasing evidence that neuronal degeneration in a number of neurodegenerative diseases is not only driven by cell-autonomous mechanisms of neuronal death, but is also promoted by non-cell autonomous processes underlying communication between neuronal and glial cells, including astrocytes [Li *et al.*, 2019; Liddelow and Barres, 2017; Siracusa *et al.*, 2019; Yamanaka and Komine, 2018].

A documented example of astrocyte involvement in neurodegeneration is provided by ALS, where an abnormal activation of astrocytes was initially suggested from the analysis of *postmortem* tissues, cerebrospinal fluids and blood samples from ALS patients [Schiffer *et al.*, 1996; Anneser *et al.*, 2004; Baron *et al.*, 2005; Poloni *et al.*, 2000]. In agreement with these observations, marked neuroinflammation was observed in the spinal cord and cranial motor nuclei of ALS transgenic mouse models [Schiffer *et al.*, 1996; Ferraiuolo *et al.*, 2007; Evans *et al.*, 2014]. Furthermore, several independent studies, based mainly on *in vitro* cultures of neural cells derived from ALS transgenic mice, provide evidence suggesting that astrocytes harbouring mutated superoxide dismutase 1 are toxic to primary, as well as pluripotent stem cell-derived, motor neurons, but not to other neuronal populations [Di Giorgio *et al.*, 2008; Marchetto *et al.*, 2008; Nagai *et al.*, 2007].

It is hypothesized that astrocytes may exert deleterious effects on motor neurons in ALS as a consequence of either gain of toxic functions or loss of supportive functions (or a combination of both). Examples of gain-of-function effects include transmission of mutated proteins such as superoxide dismutase 1, production of reactive oxygen species causing motor neuron hyperexcitability and degeneration, inhibition of autophagy mechanisms in motor neurons, and transmission of extracellular vesicle-transported toxic microRNA species, to name only a few [Fritz *at al,* 2014; Rojas *et al.*, 2015; Tripathi *et al.*, 2017; Varcianna *et al.*, 2019; Yamanaka and Komine, 2018].

Progress in understanding the roles of astrocytes in neurological diseases has been hampered by the limited availability of primary human astrocyte cultures, the limited viability of primary cultures of human astrocytes, as well as the observation that rodent and human astrocytes differ at both transcriptional and functional levels, which suggest that the rodent astrocyte models studied to date may not always represent optimal systems for understanding the pathobiology of human astrocytes [Oberheim *et al.*, 2009; Tarassishin *et al.*, 2014; Zhang *et al.*, 2016]. This situation has recently started to change thanks to the increasing availability of human iPSC-based protocols to achieve robust and reliable generation of human astrocytes [e.g., Serio *et al.*, 2013; Shaltouki *et al.*, 2013; Tcw *et al.*, 2017]. Progress in this field of research now offers previously unavailable human experimental model systems to study the biological functions of human astrocyte biology, as well as to model astrocyte pathophysiological mechanisms *in vitro.* Moreover, human iPSC-derived astrocytes have the potential to provide informative platforms for large-scale screens of new potential therapeutic compounds targeting astrocyte-mediated neurotoxic mechanisms.

In the present studies, we have described the derivation and characterization of astrocytes from a number of different human iPSC lines and have provided evidence that these cells display molecular and biological properties resembling those of physiological astrocytes. In particular, human iPSC-derived astrocytes can respond to stimulation with a pro-inflammatory combination of TNFa, IL-1a, and C1q, which are secreted by activated microglia during neuroinflammatory processes and can lead to astrocyte activation [Liddelow *et al.*, 2017], by upregulating the levels of a number of transcripts encoding molecules previously associated with a reactive astrocyte phenotype. Interestingly, elevated levels of genes previously associated with neurotoxic reactive astrocytes, such as *SERGLYCIN, SERPING1*, *GLUL* and *TNFa* [Bellaver *et al.*, 2017; Fernandes *et al.*, 2016; Liddelow *et al.*, 2017] is not accompanied by a concurrent increase in the levels of transcripts encoding intermediate filament proteins like GFAP and VIMENTIN. Increased levels of GFAP and VIMENTIN are considered a typical feature of pro-inflammatory reactive astrocytes (Clarke *et al.*, 2018; Liddelow *et al.*, 2017; Zamanian *et al.*, 2012). This situation was correlated with increased levels of *EAAT1,* encoding a glutamate transporter upregulated in reactive astrocytes with a predicted neuroprotective phenotype [Berschorner et al., 2007]. Together, these findings show a multifaceted pattern of gene expression changes in human iPSC-derived astrocytes exposed to inflammatory molecules under serum-free conditions, suggesting that human iPSC-derived astrocytes may provide informative experimental model systems to study the complex biological changes underlying reactive astrogliosis.

The derivation and characterization of human iPSC-derived astrocytes will facilitate the study of the properties and functions of different types of human astrocytes, leading to progress in our understanding of human astrocyte biology. Moreover, they will provide previously unavailable disease-relevant human experimental models to study neurological diseases. In turn, deeply phenotyped disease-relevant astrocytes may provide new experimental model systems to develop cellular assays that may contribute to early-stage drug discovery efforts benefitting neurological and mental health conditions.

## Acknowledgments

We thank Yeman Tang, Carol Chen, Rita Lo, and Tahnee Mackensen for invaluable advice and assistance. SS and GR were supported by funding from ALS Canada/Brain Canada Hudson Translational Team Grant. Additional funding was provided by the Canadian Institutes for Health Research and Fonds de la recherche en Sante-Quebec (SS).

