## Supplemental figures for "Characterization of human iPSC-derived astrocytes with potential for disease modeling and drug discovery"

### Figure S1

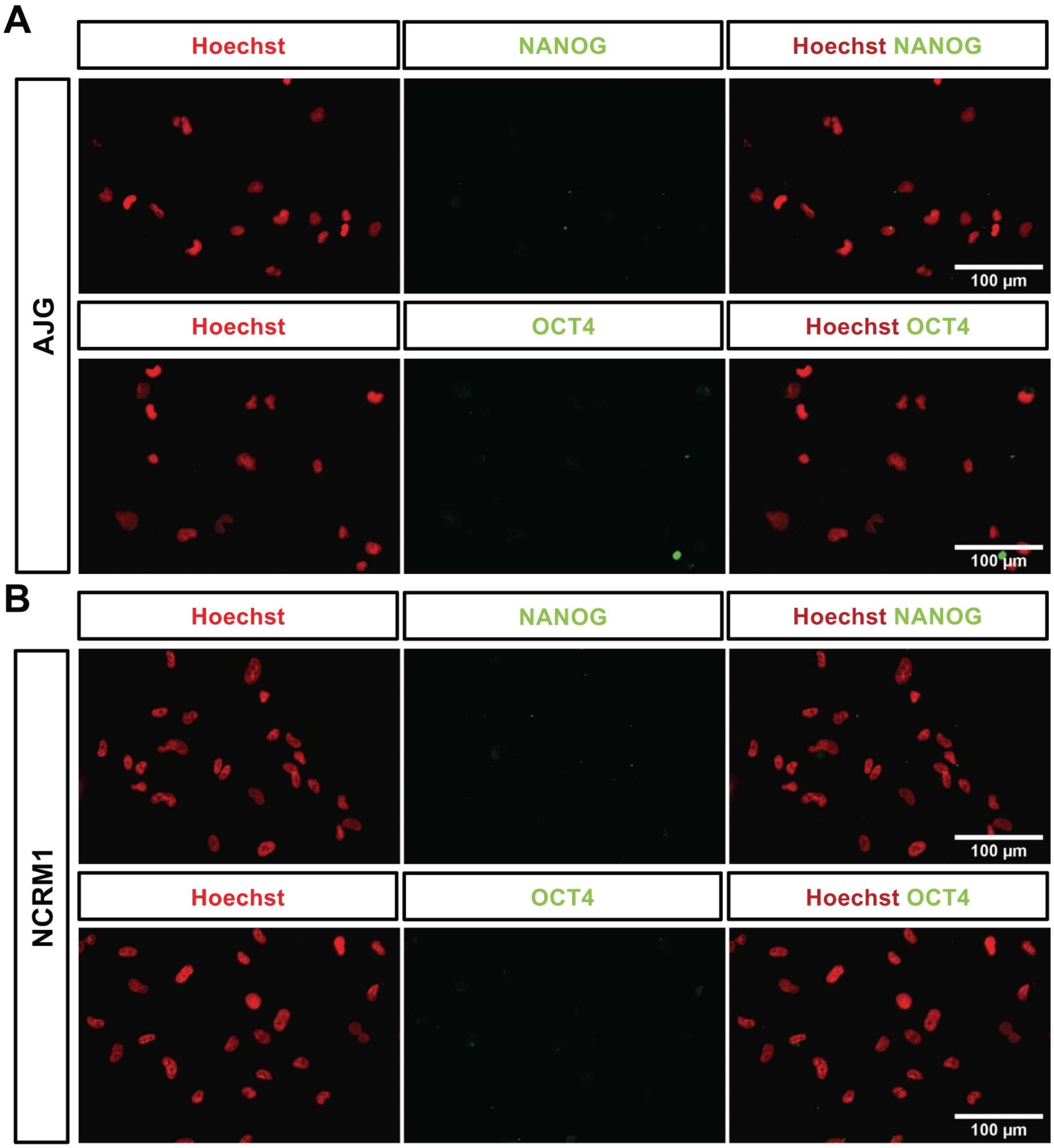


**Figure S1. Characterization of human iPSC-derived neural progenitor cells.** Representative immunofluorescence analysis of either NANOG or OCT4 expression in NPCs derived from two different iPSC lines (AJG and NCRM1). Cells were counterstained with Hoechst reagent.

**Figure S2**

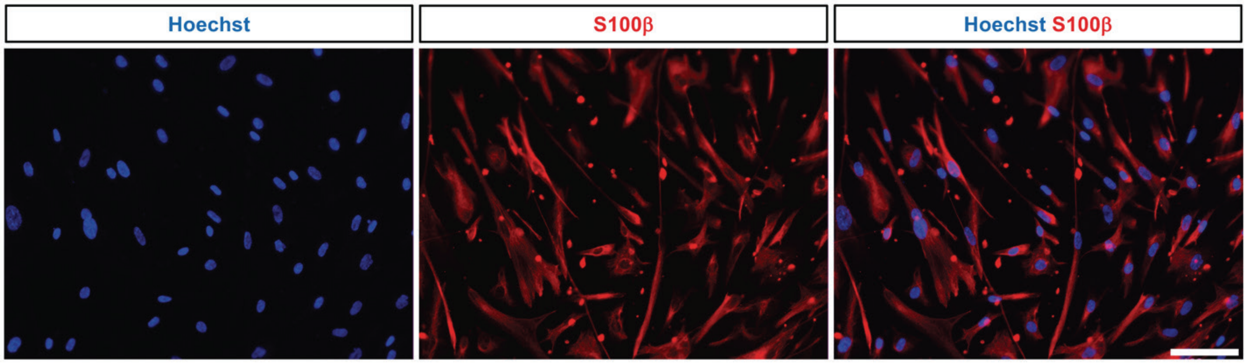


**Figure S2. Characterization of human iPSC-derived astrocytes.** Representative double-labeling immunofluorescence analysis of GFAP and S100β expression in DIV30 astrocytes derived from iPSC line AJG. Cells were counterstained with Hoechst reagent.

**Figure S3**

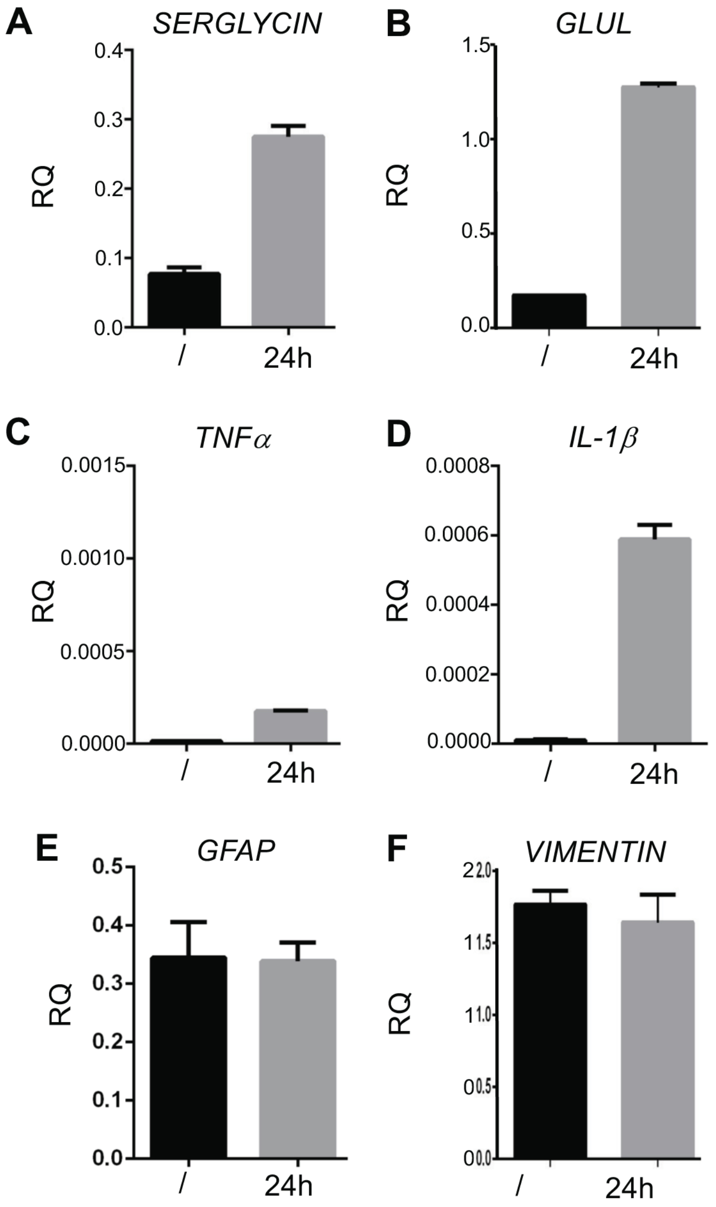


**Figure S3. Response of human iPSC-derived astrocytes to TNFα, IL-1α and C1q.** Results of real-time PCR experiments depicting mRNA levels of *SERGLYCIN*, *GLUL*, *TNFα*, *IL-1β, GFAP*, and *VIMENTIN* in DIV60 astrocytes derived from iPSC line NCRM1 not treated (/) or treated for 24h (24h) with a combination of TNFα, IL-1α and C1q. n = 2 experiments performed in triplicates.
